# Tyrosine Kinase Inhibitor Family of Drugs as Prospective Targeted Therapy for COVID-19 Based on *In Silico* And 3D-Human Vascular Lung Model Studies

**DOI:** 10.1101/2021.05.13.443955

**Authors:** Shalini Saxena, Kranti Meher, Madhuri Rotella, Subhramanyam Vangala, Satish Chandran, Nikhil Malhotra, Ratnakar Palakodeti, Sreedhara R Voleti, Uday Saxena

## Abstract

COVID-19 pandemic has ravaged the world and vaccines have been rapidly developed as preventive measures. But there is no target-based therapy which can be used if infection sets in. Remdesiver and dexamethasone were not designed to combat COVID-19 but are used clinically till better targeted therapies are available. Given this situation target based therapies that intervene in the disease pathway are urgently needed.

Since COVID-19 genesis is driven by uncontrolled inflammation/thrombosis and protein kinases are critical in mounting this response, we explored if available tyrosine kinase inhibitors (TKI’s) can be used as intervention. We profiled four TKI’s namely; Lapatinib, Dasatinib, Pazopanib and Sitravatinib which inhibit tyrosine kinases but are completely distinct in their chemical structures.

We demonstrate using *in silico* and an *in vitro* 3D-human vascular lung model which profiles anti-inflammatory and anti-thrombogenic properties that all four TKI’s are active in varying degrees. Our findings that chemically different TKI’s which share kinase inhibition as the common mechanism of action are active, strongly indicates that it’s a tyrosine kinase target-based activity and not off-target arbitrary effect. We propose that TKI’s, approved for human use and widely available, can be rapidly deployed as specific target-based therapy for COVID-19.

## 1. Introduction

Introduction of vaccines against SARS-CoV-2 spike protein has tremendously reduced the health burden of this COVID-19. Vaccine is an excellent preventive approach which can be deployed to reduce the spread of this virus. However, vaccines are not yet globally available and with potential new variants which may elude neutralization by current vaccines (escape mutants of the virus), we also urgently need a treatment for this disease once it sets in. Such drugs could save several million human lives as well as reduce hospitalization and adverse effects of the virus.

A brief proposed view of how the disease sets in and triggers inflammation and thrombosis are shown in **Figure-1**. The virus gains entry into host cells using a receptor binding domain (RBD) on its surface spike protein to bind to human host cell receptor called angiotensin receptor-2 (ACE-2). This attachment leads to internalization of the virus where it uses host machinery to replicate itself. The attachment and entry of virus into host cell results in cellular dysregulation of inflammation and thrombosis. For example, in the endothelium, it can lead to upregulation of leukocyte adhesive proteins such as Vascular Cell Adhesion Molecule-1 (VCAM-1) to attract monocytes to the site of infection to fight it by mounting this inflammatory response. It may also lead to excess secretion of pro inflammatory cytokine secretion such as TNFα and IL-1β which further exaggerate inflammation. The recruited monocytes themselves contribute to cytokine secretion causing the “cytokine storm”. The other important change during this period of inflammation is the down regulation of endothelial cell anti-thrombogenic properties. Down regulation and or secretion of thrombomodulin (which is normally functional as a cell surface protein) an anti-thrombotic endothelial surface protein, can create a pro-thrombogenic surface and increase blood clotting as seen in COVID-19 patients.

**Figure-1:**
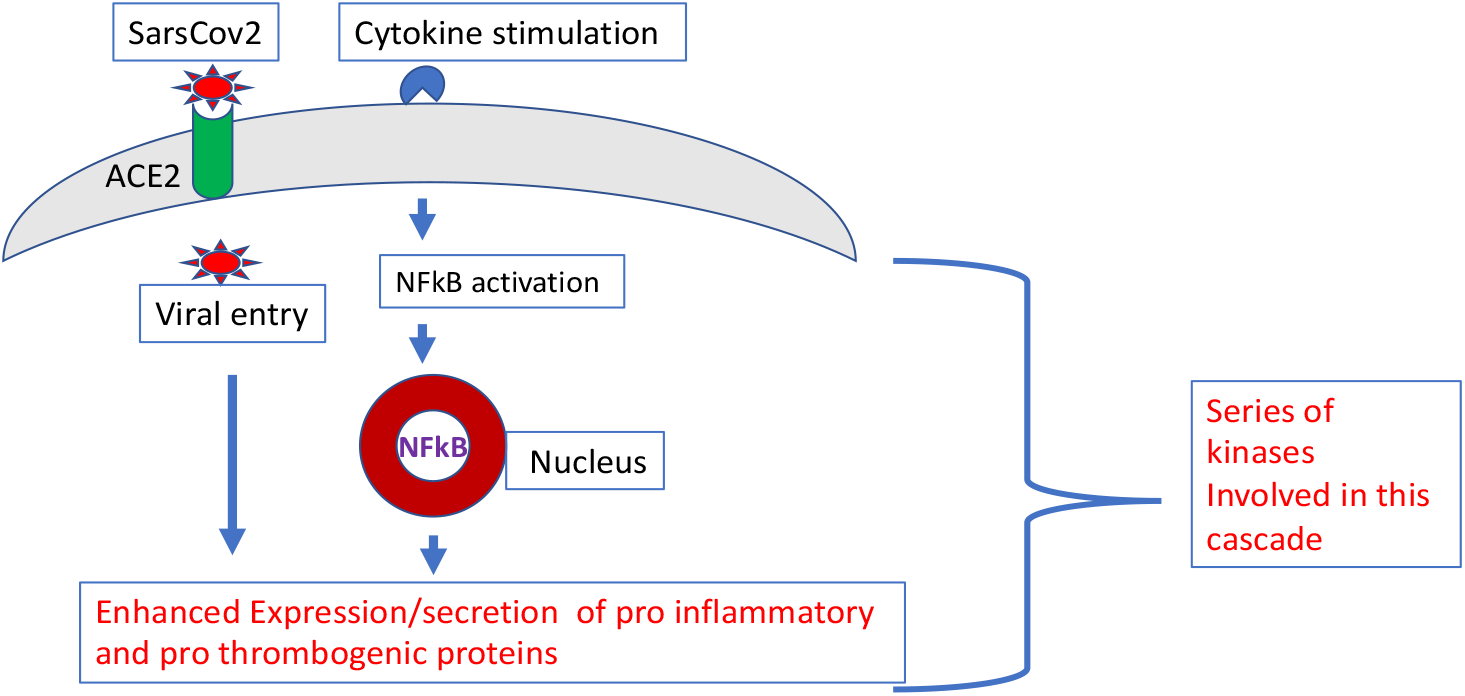
A brief view of COVID-19 triggers inflammation and thrombosis.

Underlying this pro-inflammatory and pro-thrombogenic events is a cascade of molecular steps that are regulated by tyrosine kinases. The best studied molecular pathway is the Nuclear Factor Kappa B (NFkB) pathway in inflammation is shown in Figure-1. Upon stimulation by cytokines, this NFkB protein subunit complex which normally resides in the cytoplasm is phosphorylated by kinases and migrates to the nucleus to trigger gene expression. A series of kinases are involved in the process of VCAM-1 expression and protease mediated secretion of thrombomodulin. Based on the above sequence of events, we sought to screen tyrosine kinase drugs *in silico* and using an *in vitro* 3D-human vascular lung model of disease. The purpose here was two-fold; 1) to identify TKI’s that could be repurposed for COVID-19 as a target-based disease modifying therapy and 2) to establish that TKI’s are selective targets for directly intervening in critical steps of COVID-19. Shown in **Table-1**, are the TKI drugs we choose for our studies, Lapatinib, Dasatinib, Pazopanib and Sitravatinib, which are all mainly indicated for cancers. While they share tyrosine kinase(s) as a common target, they are yet structurally different (**Figure-2**). If they all show activity in our screens, it would suggest that the effect is likely due to tyrosine kinase inhibition.

**Table-1:**
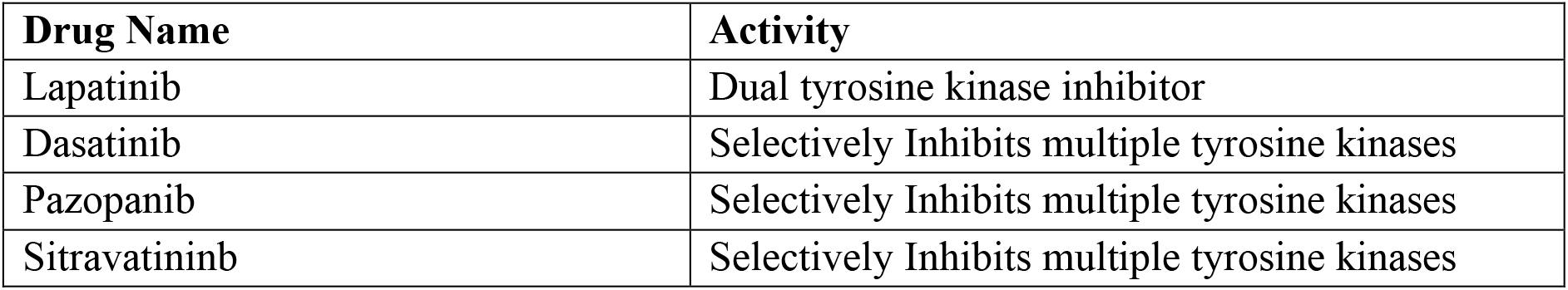
Tyrosine kinase drugs used for *in-silico and in vitro* studies.

| Drug Name | Activity |
| --- | --- |
| Lapatinib | Dual tyrosine kinase inhibitor |
| Dasatinib | Selectively Inhibits multiple tyrosine kinases |
| Pazopanib | Selectively Inhibits multiple tyrosine kinases |
| Sitravatinib | Selectively Inhibits multiple tyrosine kinases |

**Figure-2:**
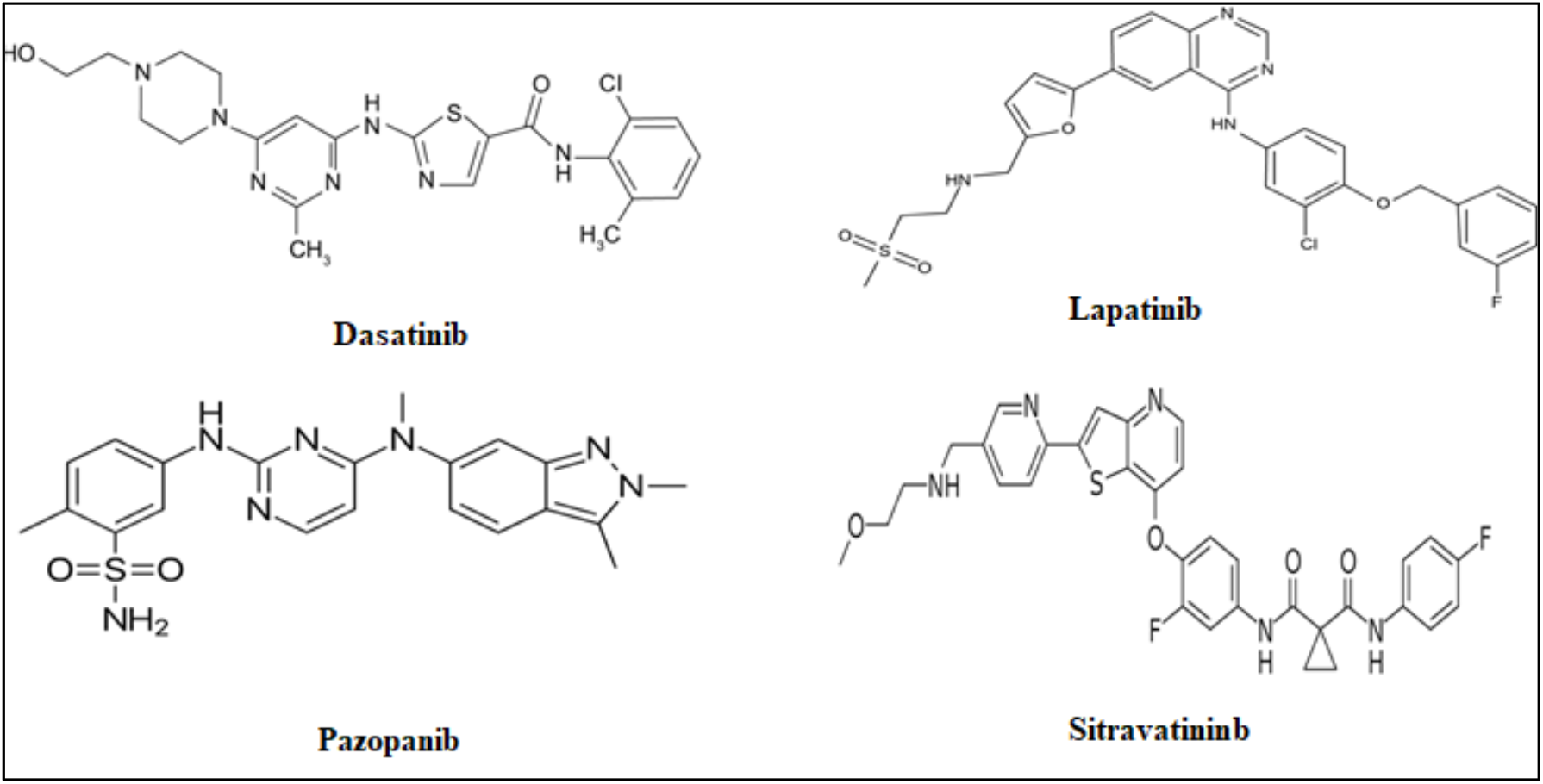
2D Structures of Tyrosine Kinase Inhibitor Drugs.

The currently used hospital paradigm for treatment of COVID-19 patients includes the use of an anti-viral drug Remdesivir, originally designed for hepatitis C, and dexamethasone which is a broad-spectrum anti-inflammatory drug, and possibly anti-cytokine *mabs* and anticoagulants. Thus, a series of drugs are needed to dampen the damage caused by the virus. If we can demonstrate in our studies that diverse TKI’s can be prospective drugs for COVID-19 it could represent the first target-based class of drugs for COVID-19. Specifically, we examined the interaction of the drugs with SARS-CoV-2’s Receptor Binding Domain (RBD) using *in silico* methods, ability to inhibit this binding in a cell-free assay, impact on inflammatory and thrombogenic properties in in vitro models. Our data suggest that TKI’s may be disease modifying drugs for COVID-19.

## 2. Material and Methods

Cresset Flare software was used for molecular docking studies against the spike protein SARS-CoV-2 (http://www.cresset-group.com/flare/) [1].

### 2.1 Ligand preparation

The 2D structures of tyrosine kinase drugs were downloaded from Drugbank database [2] and prepared using Flare software. Hydrogen atoms were added in the structure and atom force field parameterization was assigned. Further, energy minimized was done for all the drugs, nonpolar hydrogen atoms were merged, and rotatable bonds were defined. Later, ligand minimization has been carried out in Flare by Minimize tool by using Normal calculation methods. Ligands should also be prepared to assign proper bond orders and to generate the correct tautomer and/or ionization state.

### 2.2 RBD structure preparation

The RBD of spike glycoprotein SARS-CoV-2 is used for present study. This RBD binds to ACE-2 receptor on the host cell with high affinity, which makes it a key target for the novel coronavirus therapy development. The 3D structure of RBD binds to ACE-2 receptor (PDB ID: 6M0J) were downloaded from Protein Data Bank (PDB) (https://www.rcsb.org) [3]. The protein has two chain A and E, the A chain has ACE-2 receptor and E chain has RBD domain. The RBD domain has been save in to PBD format for further studies. The target protein preparation was carried out in Flare software with default settings. Missing residues, hydrogen’s and 3D protonation were carried out on the target protein. Protein minimization has been carried out in Flare by Minimize tool by using Normal calculation methods.

### 2.3 Computational analysis of binding sites

Binding site was generated Accelrys Discovery Studio visualizer 3.5 (Copyright© 2005-12, Accelrys Software Inc.) to explore potential binding sites of the RBD protein using receptor cavities tools. Based on a grid search and “eraser” algorithm, the program defines where a binding site is. The binding sites were displayed as a set of points (point count) and the volume of each of cavity was calculated as the product of the number of site points and the cube of the grid spacing. Volume of each site were calculated and further saved and exported in to Flare for advance analysis.

### 2.4 Docking with Lead Finder

The full-atom model of RBD was prepared and docking of ligands to the prepared model of RBD in the active sites was performed using Flare (Lead Finder) software (http://www.cresset-group.com/lead-finder/) by default configuration setting [4]. The energy grid box for ligand docking was set at the geometrical centre of the active site residues to span 10 Å in each direction. Two active sites were identified and in each sites centre residues were picked to define the active site grid box. Lead Finder assumes that the protein is rigid and analyses the possible conformations of the ligand by rotating functional groups along each freely rotatable bond. For each ligand pose Lead Finder determines values of the free energy of binding, the VS score, and the pose ranking score by using its three built-in scoring functions. Three different scoring functions are available from Lead Finder.

**Rank Score:** optimized to provide accurate prediction of 3D docked ligand poses.

**ΔG:** optimized to provide an accurate estimate of protein-ligand binding energy, on the assumption that the pose is correct.

**VS:** optimized to provide maximum efficiency in virtual screening experiments, with a maximum discrimination between active and inactive compounds in virtual screening experiments.

The selected drugs were docked in the RBD domain of S-protein SARS-CoV-2 by using Cresset Flare Docking software with accurate and slow mode. The best poses were generated and visualized in pose viewer and 3D images stored in storyboard.

### 2.5 RBD-ACE-2 biochemical assay

The RBD-ACE2 assay was performed using SARS-CoV-2 sVNT ready to use kit sold by Genscript Inc, USA. It is which is a competition ELISA, mirroring the viral neutralization process. In the first step, a mixture of horse radish peroxidase-RBD (HRP-RBD) and controls/drugs (drugs at the concentrations of 50μM, 100μM and 200μM) were incubated at 37°C for an hour to allow the binding of drugs to HRP-RBD. Following the incubation, these mixtures were added to a capture plate, which was a 96 well microplate coated with human ACE-2 receptor to permit the binding of any free HRP-RBD. After incubating the microplate at 37°C for an hour, the plate was washed four times using wash buffer in order to remove any unbound HRP-RBD drug complexes. Washing step was followed by addition of a colour substrate; tetramethyleneblue (TMB), which yields blue color. The reaction was allowed to run for 15 minutes followed by the quenching using stop solution turning the colour from blue to yellow. This final solution was read at absorbance 450 nm. The absorbance of the sample is inversely proportional to the inhibition of RBD’s binding to ACE-2 by the drug. Data is presented as percent inhibition relative to untreated wells.

### 2.6 Interleukin −1 β (IL-1β) secretion assay

Human IL-1**β** ELISA was performed to detect the presence of IL-1**β** which is a key mediator of inflammatory response. The ELISA was performed using Human IL-1**β** high sensitivity kit from Invitrogen Inc, USA. The sample of ELISA used in the present study consist of cell culture media (referred to as sample hereafter) collected after treating the cells with drugs at 100μM. Samples were added to the microplate pre-coated with human IL-1**β** antibody which captures the IL-1 **β** present in the samples. A secondary anti-human IL-1**β** antibody conjugated to biotin was added to the plate. Following an overnight incubation, microplate was washed six times using wash buffer in order to remove any unbound biotin conjugated anti-human IL-1**β** antibody. Streptavidin-HRP was then added which binds to biotin conjugated antibody and the plate was incubated at room temperature on a shaker for an hour. After the incubation, plate was washed again following the same process as the previous wash step and an amplification reagent-I was added to the wells. Following the incubation of 15 minutes and a wash, amplification reagent-II was added. After incubation of half an hour in dark and a wash step later, a substrate solution was added turning the colour to blue. After 15-20 minutes, the reaction was terminated using a stop solution (turning the colour from blue to yellow). This final solution was read at 450nm. The OD of the sample is directly proportional to the amount of human IL-1**β** present in it.

### 2.7 Human Thrombodulin/BDCA-3 Immunoassay

Thrombomodulin is a transmembrane glycoprotein expressed by a variety of cells, including endothelial cells. Human Thrombodulin ELISA was performed using a ready to use sandwich ELISA kit by R and D systems. Samples (media collected from our 3D-vascular lung system) were added to a microplate precoated with monoclonal antibody specific for human thrombodulin. This was followed by an incubation of two hours (allowing any thrombodulin present in the sample to bind to the monoclonal antibody), the plate was washed with wash buffer four times. After washing an enzyme-linked monoclonal antibody specific for human thrombodulin was added to the plate. Another wash step was performed in order to remove any unbound antibody-enzyme reagent after completion of two hours of incubation. A colour substrate was then added turning the colour to blue. Reaction was quenched after 15-20 minutes using a stop solution which turned the colour from blue to yellow. Absorbance was read at 450 with wavelength correction set to 540 nm or 570 nm. Data is presented as amount of thrombomodulin in the media.

### 2.8 3D-Vascular Lung Model

#### 2.8.1 Cell culture

For the 3D vascular lung model three types of cell were used:

1. A549 cellss, lung epithelial cells, were grown at 37° C in the growth medium DMEM (HIMEDIA #Cat No-AT007) supplemented with 10% (v/v) fetal bovine serum under the atmosphere containing 5% CO2. Cells were subcultured after reaching 80-90% confluence.
2. Human umbilical vein endothelial cells (HUVEC) cells were grown at 37° C in the Endothelial cell basal medium-2 (LONZA #Cat No-CC-3156 and CC-4176) under the atmosphere containing 5% CO2. Cells were subcultured after reaching 80-90% confluence.
3. HL60, human monocytic cells were grown at 37° C in the growth medium RPMI (HIMEDIA #Cat No-AT028) supplemented with 10% (v/v) fetal bovine serum under the atmosphere containing 5% CO2. Cells were subcultured after reaching 80% - 90% confluence.

#### 2.8.2 3D-bioprinting

In the 3D-vascular lung model four layers were bioprinted using a robotic computerized 3D printer with specific flow velocity and volume dispensation in 96 well plates.. The first layer to be printed was collagen layer, (30μl of Rat tail collagen). It added to each well in 96 well plate and incubated for 1 hour in CO2 incubator at 37°C. After incubation, once the collagen is solidified A549 cells flask which has reached 80-90% confluence were trypsinized and cells were counted with the help of haemocytometer. Cell suspension was loaded into bioprinter syringe and 5×10^3^ A549 cells were printed in each well of 96 well plate and the cells were incubated for 48 hours. After 48 hours of incubation, previous A549 media was removed carefully and again another layer of 30μl of Rat tail collagen coating solution was printed in each well in 96 well plate and incubated for 1 hour in CO2 incubator at 37°C. After incubation, once the collagen is solidified HUVEC cells flask which has reached 80-90% confluence were trypsinized. Cell suspension was loaded into syringe and 5×10^3^ HUVEC cells were printed in each well of 96 well plate with help of 3D bioprinter and the cells were incubated for 48 hours. After 48 hours of incubation the media was removed and the final selected drugs from virtual screening were added with LPS at 0.5 μg/ml and without LPS at the desired concentration and cells incubated overnight. Next day the drugs were removed and the cells were washed once with media. Endothelial cells were fixed with 4% paraformaldehyde for 3 minutes at 40°C. After fixing with paraformaldehyde, the cells were again washed with media. MTT stained HL60 monocytic cells 1×10^4^ cells were added to each well in 96 well plate and incubated for one hour and the wells were then washed twice with the media. Pictures were taken of all the wells and the bound HL 60 cells were counted with the help of ImageJ, an open access software platform.

## 3. Results and Discussion

### 3.1 Identification of active sites in Receptor Binding Domain

In *silico* approach helped us to screen new drug targeting RBD domain which plays a pivotal role in viral-host attachment. One such avenue of research is to find repurposed drug(s) that can attach to residues at the site of binding of the RBD to the ACE-2, thus disrupting the protein-protein interactions. In the current study, we identified several potential binding sites for molecules that can occupy such druggable pockets so as to inhibit virus-ACE-2 binding *in vitro*. The X-ray crystal structure of the RBD was used to identify *3 possibly druggable pockets* where drugs possibly could bind. The active site volume and binding surface area of three pocket is representation in **Table-2** [5]. Site-1 has the volume of 143.7 **Å^3^** & site-2 has the active site volume of 109.4 **Å^3^**, the last site-3 has active site volume of 87.5 (**Å^3^**). Site-3 was too small to accommodate ligands, so it could not be a potential pocket. Based on size of active site cavity site-1 & site-2 has been selected for further docking studies.

**Table-2:** Active sites of RBD

| Site | Binding surface area (Å<sup>2</sup>) | Active site volume (Å<sup>3</sup>) |
| --- | --- | --- |
| S1-1 | 673.6 | 143.7 |
| S1-2 | 668.3 | 109.4 |
| S1-3 | - | 87.5 |

### 3.2 Docking with Lead Finder (LF)

After the identification of active sites, all the tyrosine kinase drugs were subjected to docking using LF and the predicted binding poses were analysed. In the present study, we used Structure-Based Virtual Screening (SBVS) which uses molecular docking techniques where all the selected drugs were docked in the RBD domain of spike protein. The compounds are scored based on the predicted interactions with the target protein and those with the top scores (hits) are selected for further analysis. All the selected drugs were docked in the spike protein SARS-CoV-2 (PDB ID: 6M0J, E-chain) by using Flare module of Cresset software in “slow & accurate mode”. All the selected drugs four kinase drugs were showing good binding affinity towards the RBD domain of spike protein in both the selected active sites. Detailed binding analysis of selected drugs towards active site of spike protein SARS-CoV-2 is accounted below. Interaction analysis of drugs with spike protein SARS-CoV-2 (RBD) were carried out to identify the compound having highest binding affinity with target protein.

Active site-1 composed of Arg454, Phe456, Arg457, Lys458, Glu465, Arg466, Asp467, Ile468, Ser469, Glu471, Thr473, Gln474 and Pro491 amino acid residues, while the site-2 composed of Leu335, Cys336, Pro337, Phe338, Gly339, Trp436, Phe342, Asn343, Val362, Ala363, Asp364, Val367, Leu368, Ser371, Ser373 and Phe374 amino acid residues, as shown in **Figure-3**. Active site-1 is more hydrophobic in nature as compare to active site-2.

**Figure-3:**
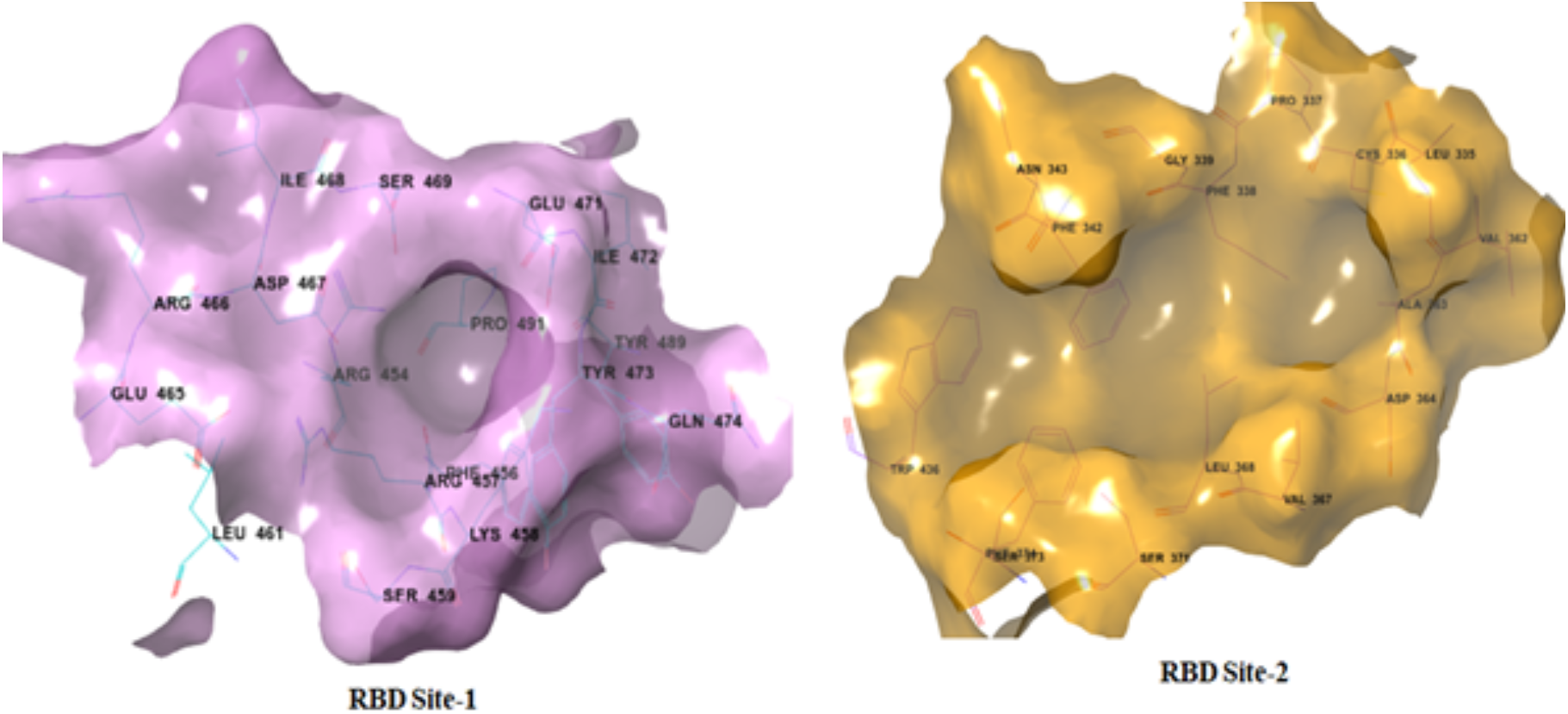
Active sites of RBD domain of Site-1 protein

The rank score was found in the range of −11.1 to 12.5 in site-1, while in site-2 the rank score range was found to be −8.8 to −11.9. The ΔG provide an accurate estimate of protein-ligand binding energy, on the assumption that the pose is correct. The ΔG of selected tyrosine kinase drugs were in the range of −7.9 to 10.4 in site-1 and −7.8 to −9.8 in site-2. From the detailed docking analysis, it is observed that these drugs have good LF rank score and ΔG in both the active sites (**Table-3**). It is found that, these four drugs have formed strong H-bond contacts with more than two to three amino acid residues along with various hydrophobic interactions in spike protein resulting in increased binding affinity with target protein.

**Table-3:** Docking score and binding energies of selected tyrosine kinase drugs for RBD domain of spike protein in site-1& site-2.

|  | RBD Site-1 |  | RBD Site-2 |  |
| --- | --- | --- | --- | --- |
| Drug Name | Rank score | $\Delta G$ | Rank score | $\Delta G$ |
| Dasatinib | -11.1 | -10.4 | -10.5 | -9.8 |
| Lapatinib | -11.6 | -8.8 | -10.8 | -7.8 |
| Pazopanib | -11.9 | -7.9 | -11.9 | -7.8 |
| Sitravatinib | -12.5 | -8.5 | -11.9 | -9.7 |

### 3.3 Protein-Ligand Interaction Analysis in RBD Domain

From the detailed docking analysis, it is observed that all four drugs (**Dasatinib**, **Lapatinib, Pazopanib, Sitravatinib**) have good LF rank score and ΔG in both the active sites. The first screened drug **Dasatinib** is a tyrosine kinase inhibitor used for the treatment of lymphoblastic or chronic myeloid leukemia (CML). **Lapatinib** is an antineoplastic agent and tyrosine kinase inhibitor used for the treatment of advanced or metastatic HER-negative breast cancer in patients who received prior chemotherapeutic treatments. Both these drugs are having good binding affinity towards the RBD domain of spike protein. In site-1, **Dasatinib** is making three hydrogen bonding interactions with Lys458, Glu465, Asp467. Further, it is involved in various hydrophobic interaction with Phe456 Leu461, Ser469, Ile472, Tyr473, Cys480 and Gly482 amino acid residues. Interaction analysis in active site-2 revealed that it has two hydrogen bonding interactions with Asn343 and Asp364 amino acid residues. (**Figure-4**). **Lapatinib** is having rank scores of −11.6 and −10.8 in site-1 and site-2, respectively, and suggests strong hydrogen bonding interactions with four active site residues (Lys458, Ser459, Glu471, Gln474) of site-1. Analysis of same drug in the active site-2 revealed that it has hydrogen bond interaction with Asn334 along with strong *pi-pi* interaction with Phe342 and Trp436, as shown in (**Figure-5**).

**Figure-4:**
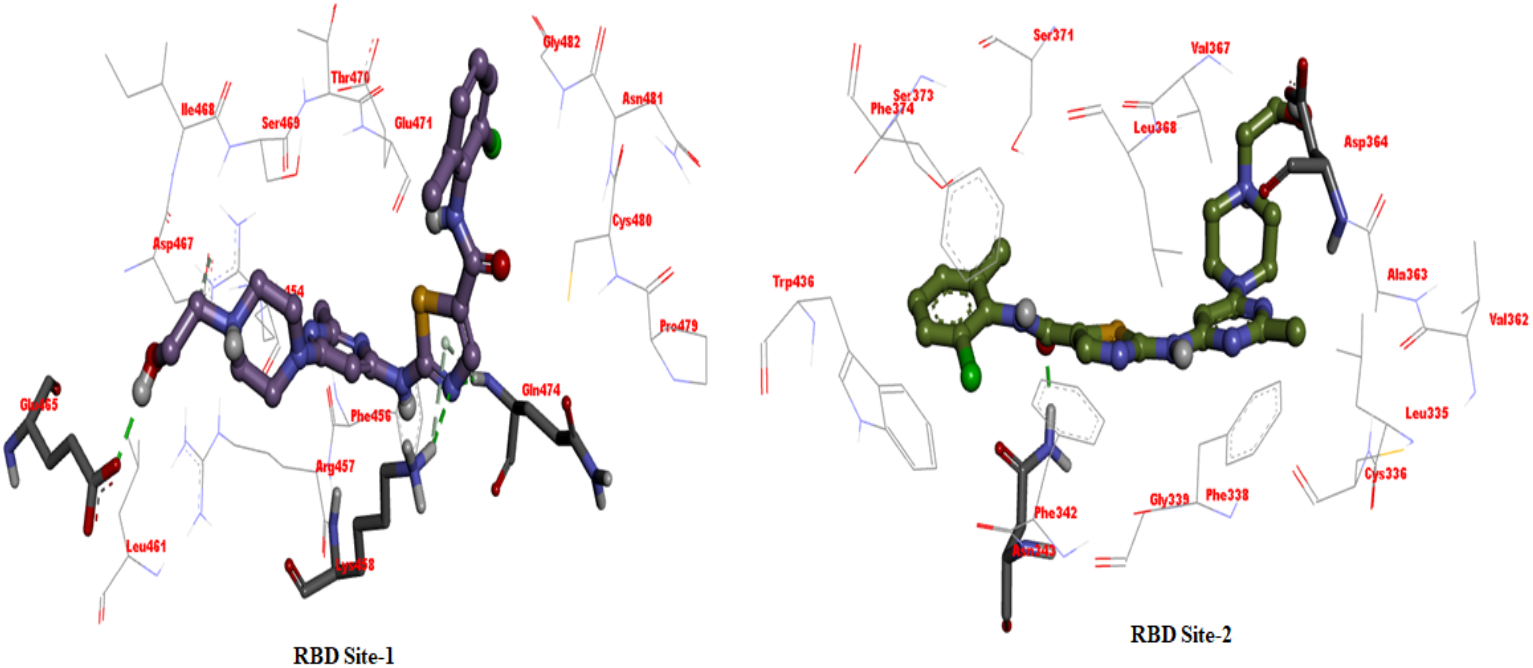
Proposed binding orientation of the drug **Dasatinib** within the active site of the RBD site-1 and site-2.

**Figure-5:**
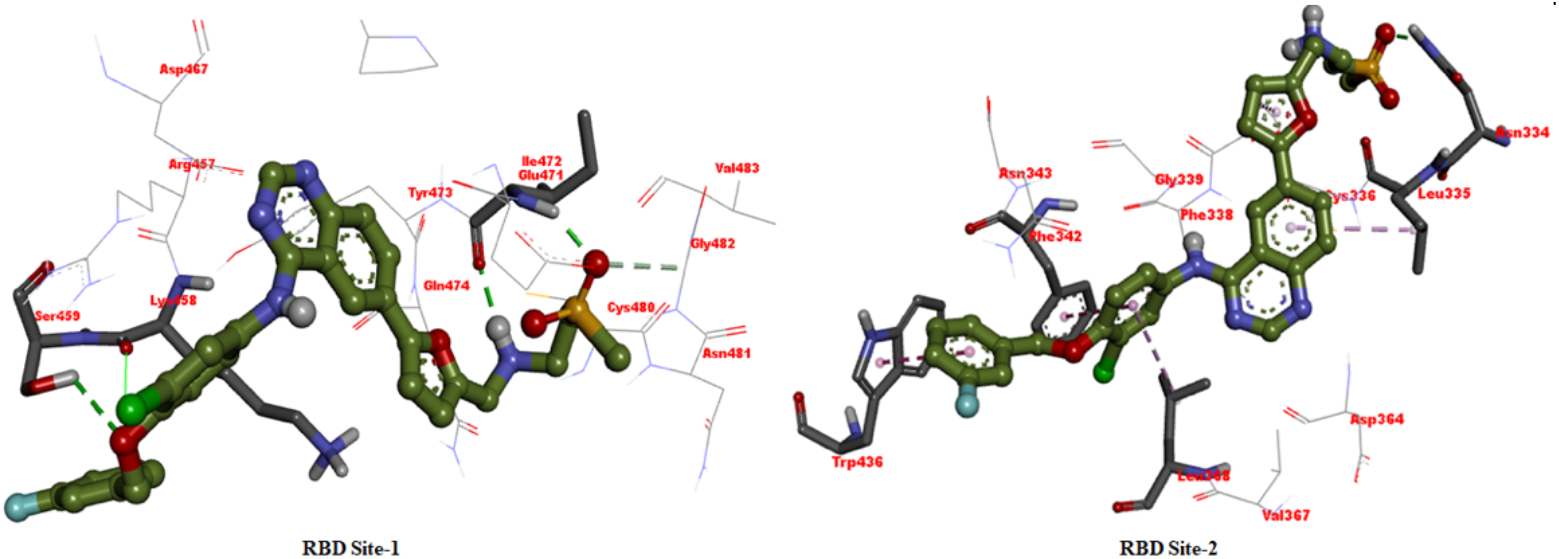
Proposed binding orientation of the drug **Lapatinib** within the active site of the RBD site-1 and site-2.

**Pazopanib** is a small molecule inhibitor of multiple protein tyrosine kinases with potential antineoplastic activity used in the treatment of advanced renal cell cancer and advanced soft tissue sarcoma in patients with prior chemotherapy. In site-1, **Pazopanib** seem to making four hydrogen bonding interactions with Arg454, Lys458, Asp467, Ser469 amino acid residues, while in site-2 it is making three hydrogen bonding interactions with Leu335, Phe342, Asn343. Apart from this some non-bonded interactions were also observed with Val367, Leu368 and *pi-pi* interactions with Phe373 and Phe374 amino acid residues (**Figure-6**).

**Figure-6:**
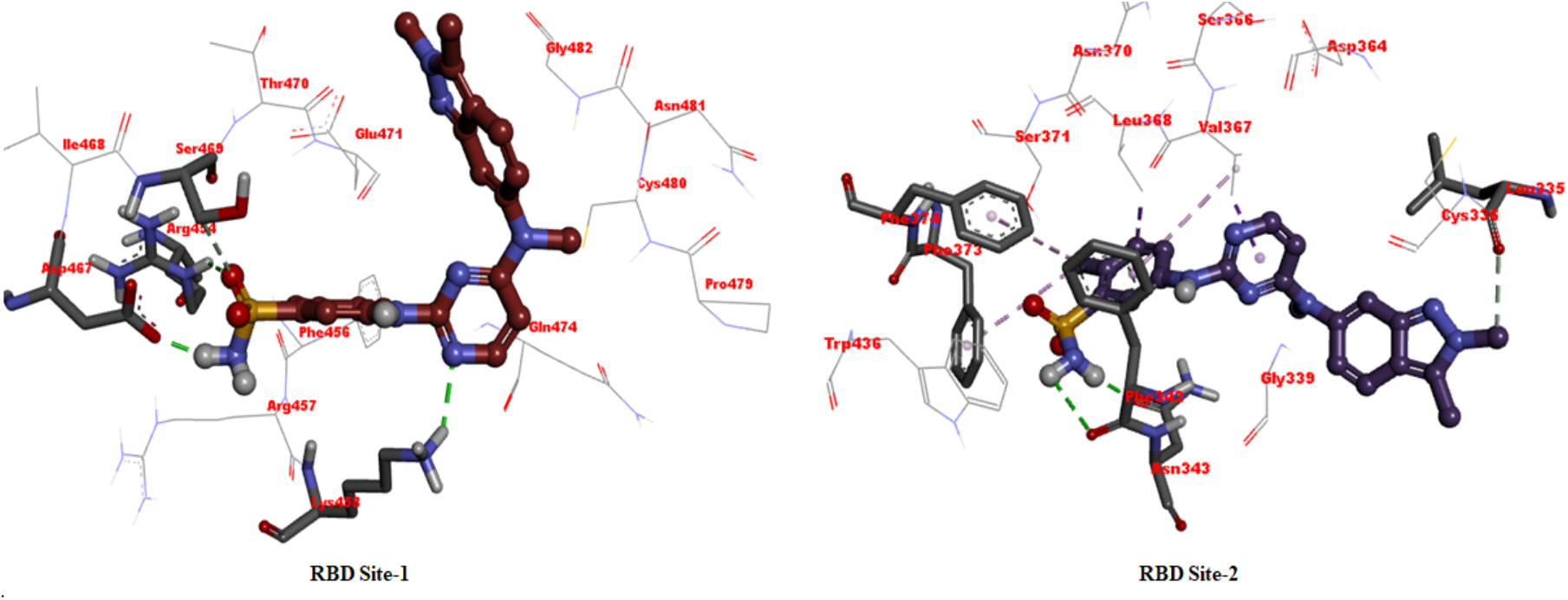
Proposed binding orientation of the drug **Pazopanib** within the active site of the RBD site-1 and site-2.

The last screened drug **Sitravatinib** is under investigation in clinical trial NCT03680521 and is also showing good binding affinity towards RBD having a rank score of −12.5 in site-1 and −11.9 in site-2, respectively. (**Figure-7**). In site-1, **Sitravatinib** showed conventional hydrogen boding interactions with Cys480, Phe456, Arg457, Lys458, Asp467 and Gln474 amino acids, while in site-2, it was making strong hydrogen bonding interactions with Trp436, Asn343, Ser371.

**Figure-7:**
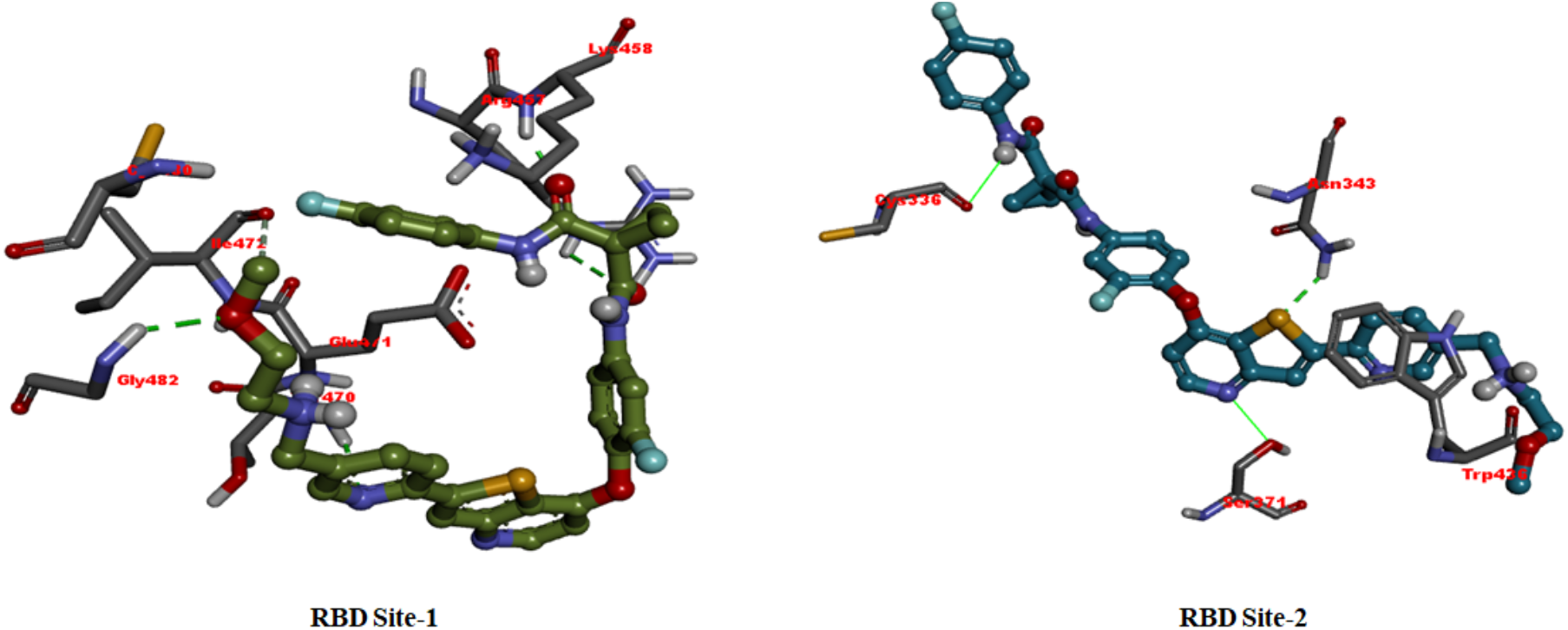
Proposed binding orientation of **Sitravatinib** within the active site of the RBD site-1 and site-2.

### 3.4 Inhibition of binding of RBD to ACE-2

To verify the *in silico* data from above that the TKI’s may block the binding to SARS-CoV-2 spike protein RBD to human ACE-2, we performed an ELISA based binding assay which measures the direct binding of RBD to ACE-2. As shown in **Figure-8**, all of the four TKI’s were able to block the binding and the maximal inhibition obtained by all was > 80% of control (no drug added). Dasatinib and Lapatinib appear to be moderately better than Pazopanib and Sitravatinib. These data corroborate the *in silico* data, and, demonstrate the ability to potentially block viral entry into host cells.

**Figure-8:**
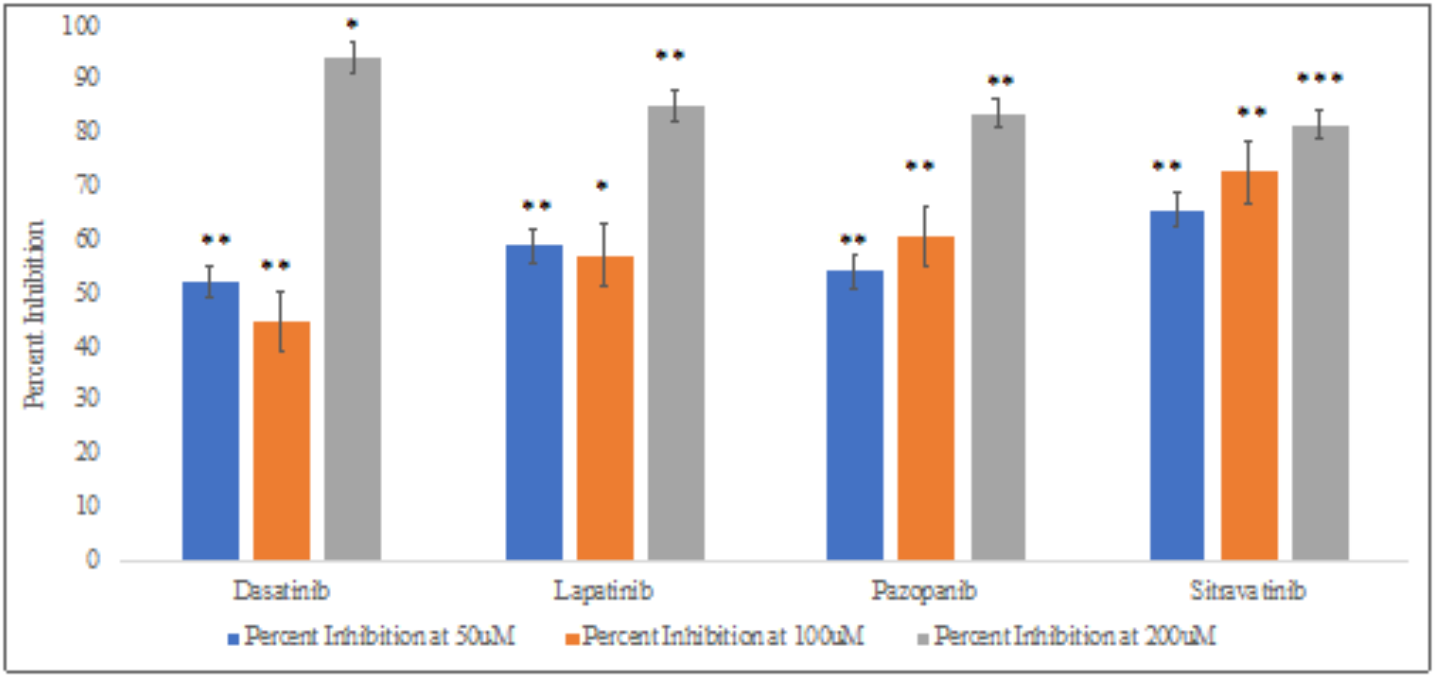
Percent Inhibition of RBD binding to ACE-2 by different drugs at different concentrations of 50μM, 100μM and 200μM. *** P<0.001; ** P<0.01; *P<0.05. Student’s t-tests were performed to compare between the percent inhibition shown by control and drugs.

**Figure-9:**
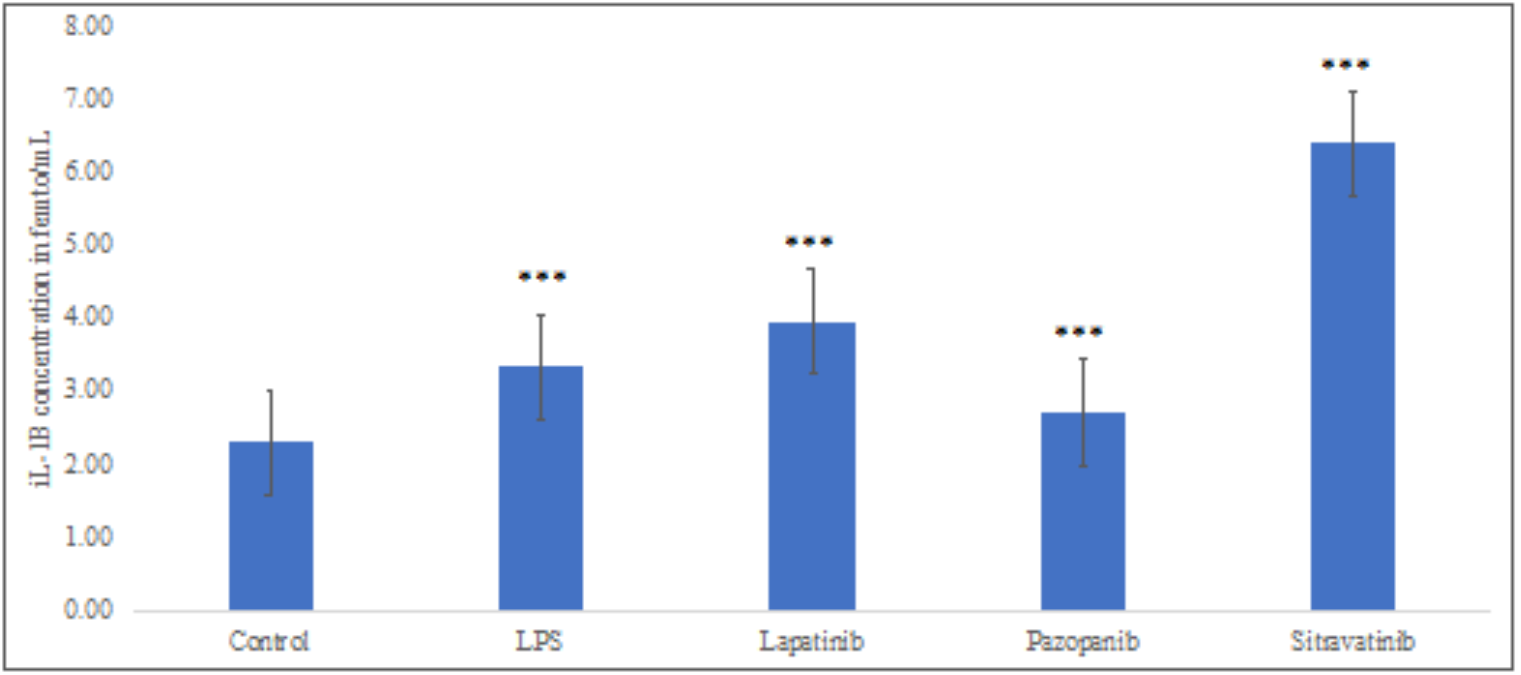
Effect of different drugs on IL-1β secretion (all drugs added at 100μM). *** P<0.001; ** P<0.01; *P<0.05. Student’s t-tests were performed to compare IL-1β secretion between control and LPS treated cells; and between LPS treated cells (acting as a control) and different drugs treated cells.

### 3.5 Inhibition of IL-1β secretion

We then tested the ability of these drugs to inhibit the secretion of IL-1β, believed to be an important cytokine in COVID-19 genesis. We used lipopolysaccharide (LPS) as a stimulus to induce IL-1 β secretion in the presence or absence of drugs added. Pazopanib was able to reduce the secretion of IL-1β significantly but the other TKI’s were not very effective.

### 3.6 Inhibition of thrombomodulin secretion

Increased thrombodulin secretion has been correlated with COVID-19 severity in patients [6]. Secreted thrombodulin cannot serve its normal anti-thrombotic function which requires it to be on endothelial cell surface. We studied whether the TKI’s can inhibit LPS induced secretion of thrombodulin. **Figure-10** shows that all four of the drugs studied herein have demonstrated reduced thrombodulin secretion, with **Dasatinib** being the most effective. Thus, these drugs have the potential to be ant-thrombogenic in terms of thrombodulin secretion.

**Figure-10:**
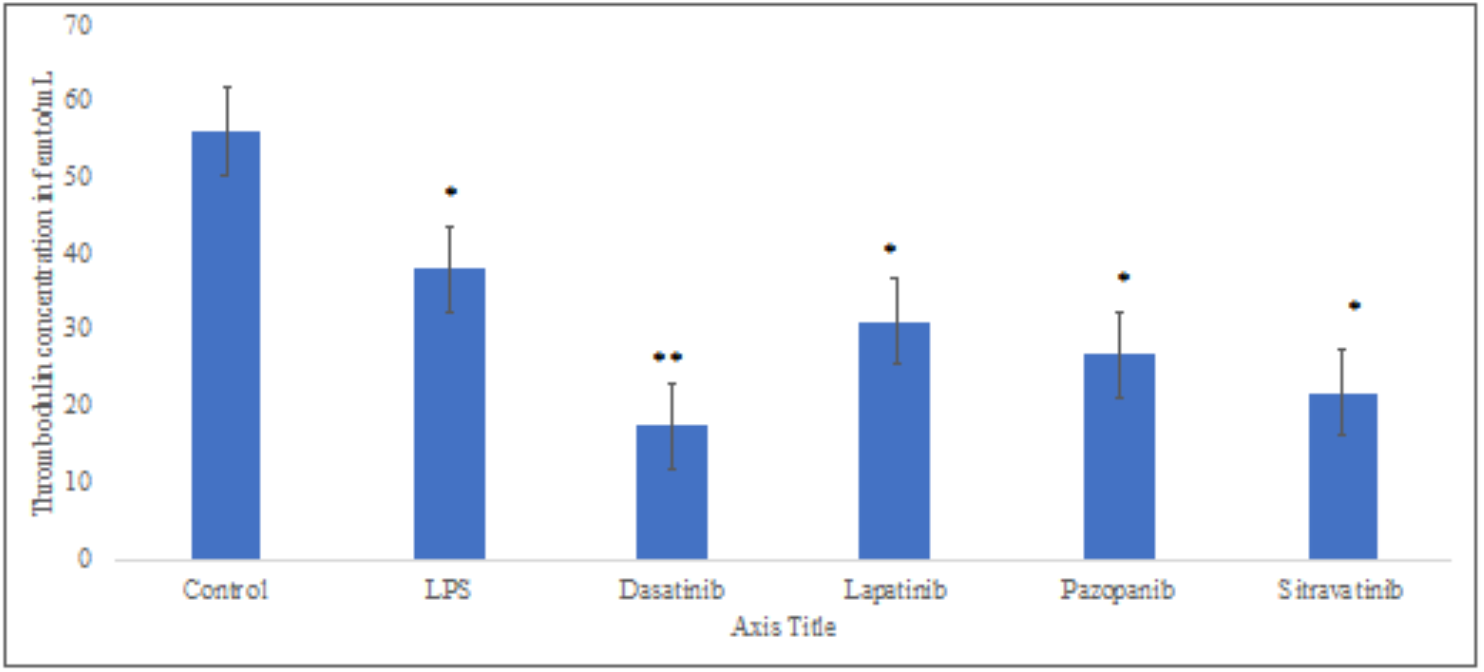
Effect of different drugs on thrombodulin secretion (all drugs added at 100μM). *** P<0.001; ** P<0.01; *P<0.05. Student’s t-tests were performed to compare thrombodulin secretion between control and LPS treated cells; and LPS treated cells and different drugs treated cells.

### 3.7 Inhibition of monocyte adhesion

We next studied the effect of TKI’s on LPS induced adhesion of monocytes to the vascular endothelium in the *in vitro* 3D model. Adhesion of monocytes is the first step in inflammation that occurs during viral infections. Once the monocytes bind, they then transmigrate into the lung tissue and start secreting cytokines. The drug which had the best inhibitory effect on adhesion (**Figure-11**) was Lapatinib which completely blunted the adhesion induced by LPS. The other drugs were less effective but showed a trend towards reduction in increase compared to the untreated cells (control) treated with LPS alone.

**Figure-11:**
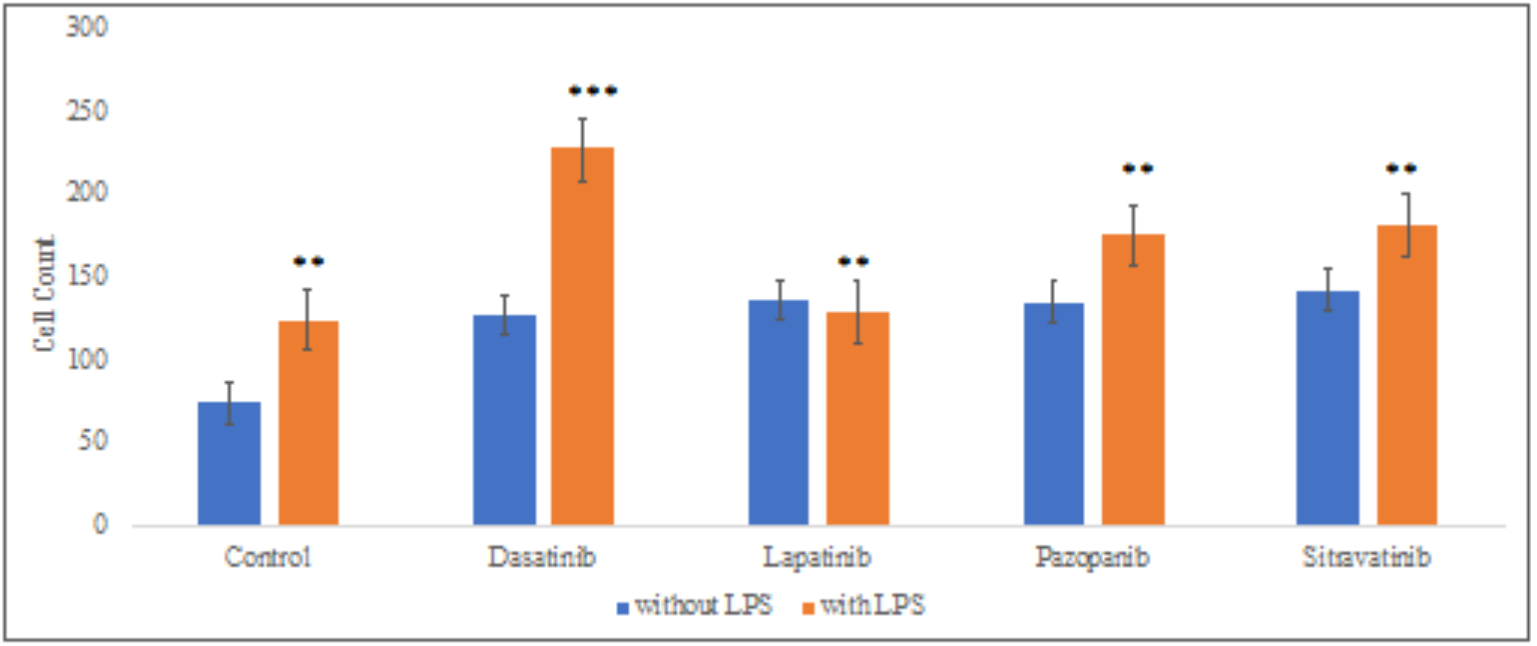
Effect of different drugs on Monocyte adhesion in 3D Vascular Lung Model. *** P<0.001; ** P<0.01; *P<0.05. Student’s t-tests were performed to compare between the control and drugs after treatment with LPS

## 4. Conclusion

The focus of this work was to evaluate if TKI’s can be repurposed for mitigation of COVID-19 through blocking the binding to RBD domain with ACE-2 as well as other antiinflammatory and anti-thrombogenic properties. The critical outcome from our studies is the finding that structurally diverse TKI’s were all able to modulate RBD binding to ACE-2 and demonstrate inflammatory and thrombogenic steps that are involved in COVID-19. The ability of TKI’s to modulate inflammatory and thrombogenic events is especially meaningful in the context of COVID-19. From human studies it is obvious that these two events are the hallmark of this viral disease and probably cause most of the damage seen in lungs, kidneys, and the brain. We are most excited about the ability of TKIs to inhibit the secretion of thrombodulin, an established circulating biomarker which correlates with disease severity in COVID-19 patients [6]. If these TKIs are able to produce this effect, as seen in from in vitro studies, in a clinical setting it will no doubt be very important for their therapeutic ability.

**Lapatinib** and **Dasatinib** have been shown to be effective in blocking SARS-Cov-2 replication by others using in vitro cell-based assays, thus strengthening our observations of their use in COVID-19 [7–9]. Furthermore, **Baricitinib**, an inhibitor of JAK tyrosine kinase, has received emergency use authorization by the US FDA to be used as treatment for COVID-19 [10]. Collectively these data together with our data shown here establish the utility of TKI’s as a first-in-class target based therapy. The more drug options that are available to treat this disease the better it is for patients. Considering the beneficial activity demonstrated by **Lapatinib** in multiple steps in our 3D human vascular lung model, we propose utilization of this drug for treatment of COVID-19 either as monotherapy or in combination with drugs currently in use for mitigation of this disease.

## References

1. Flare, version, Cresset®, Litlington, Cambridgeshire, UK; http://www.cresset-group.com/flare/

2. Wishart, D.S., Knox, C., Guo, A.C., Shrivastava, S., Hassanali, M., Stothard, P., Chang, Z. and Woolsey, J. DrugBank: a comprehensive resource for *in silico* drug discovery and exploration. Nucleic acids research, 34, 668–672. 2006.

3. Lan, J., Ge, J., Yu, J., Shan, S., Zhou, H., Fan, S., and Wang, X. Structure of the SARS-CoV-2 spike receptor-binding domain bound to the ACE-2 receptor. Nature, 581(7807), 215–220. 2020.

4. Lead Finder, version, BioMolTech®, Toronto, Ontario, Canada; http://www.cresset-group.com/lead-finder/

5. Saxena, S., Meher, K., Rotella, M., Vangala, S., Chandran, S., Malhotra, N. Saxena, U. *In silico* and *in vitro* Demonstration of Homoharrintonine Antagonism of RBD-ACE-2 Binding and its Anti-inflammatory and anti-thrombogenic Properties in a 3D human vascular lung model. bioRxiv. doi: 10.1101/2021.05.02.442384. 2021.

6. Goshua, G., Pine, A.B., Meizlish, M.L., Chang, C.H., Zhang, H., Bahel, P., Baluha, A., Bar, N., Bona, R.D., Burns, A.J. and Cruz, C.S.D. Endotheliopathy in COVID-19-associated coagulopathy: Evidence from a single-centre, cross-sectional study. The Lancet Haematology, 7(8), 575–582. 2020.

7. Weisberg, E., Parent, A., Yang, P.L., Sattler, M., Liu, Q., Liu, Q., Wang, J., Meng, C., Buhrlage, S.J., Gray, N. and Griffin, J.D. Repurposing of kinase inhibitors for treatment of COVID-19. Pharmaceutical research, 37(9), 1–29. 2020.

8. O’Donovan, S.M., Imami, A., Eby, H., Henkel, N.D., Creeden, J.F., Asah, S., Zhang, X., Wu, X., Alnafisah, R., Taylor, R.T. and Reigle, J. Identification of candidate repurposable drugs to combat COVID-19 using a signature-based approach. Scientific reports, 11(1), 1–12. 2021.

9. Raymonda, M.H., Ciesla, J.H., Monaghan, M., Leach, J., Asantewaa, G., Smorodintsev-Schiller, L.A., Lutz, M.M., Schafer, X.L., Takimoto, T., Dewhurst, S. and Munger, J. Pharmacologic profiling reveals lapatinib as a novel antiviral against SARS-CoV-2 *in vitro*. bioRxiv. doi: 10.1101/2020.11.25.398859. 2020.

10. Kalil, A.C., Patterson, T.F., Mehta, A.K., Tomashek, K.M., Wolfe, C.R., Ghazaryan, V., Marconi, V.C., Ruiz-Palacios, G.M., Hsieh, L., Kline, S. and Tapson, V. Baricitinib plus Remdesivir for hospitalized adults with Covid-19. New England Journal of Medicine, 384(9), 795–807. 2021.

